# *MYB5a/NEGAN* activates petal anthocyanin pigmentation and shapes the MBW regulatory network in *Mimulus luteus* var. *variegatus*

**DOI:** 10.1101/2020.04.09.030536

**Authors:** Xingyu Zheng, Kuenzang Om, Kimmy A. Stanton, Daniel Thomas, Philip A. Cheng, Allison Eggert, Yao-Wu Yuan, Joshua R. Puzey, Arielle M. Cooley

**Author notes:** School of Biological Sciences, Washington State University, Pullman, WA, U.S.A. Corresponding author for transcriptome analysis, Corresponding author for sequence analysis, RT-PCR, and transgenics.

## Abstract

Much of the visual diversity of angiosperms is due to the frequent evolution of novel pigmentation patterns in flowers. The gene network responsible for anthocyanin pigmentation, in particular, has become a model for investigating how genetic changes give rise to phenotypic innovation. In the monkeyflower genus *Mimulus*, an evolutionarily recent gain of petal lobe anthocyanin pigmentation in *M. luteus* var. *variegatus* was previously mapped to genomic region *pla2*. Here, we use DNA sequence analysis and spatiotemporal patterns of gene expression to identify *MYB5a* - homologous to the *NEGAN* transcriptional activator from *M. lewisii* - as a likely candidate gene within the *pla2* region. Transgenic manipulation of gene expression confirms that *MYB5a* is both necessary and sufficient for petal lobe anthocyanin pigmentation. The deployment of *MYB5a/NEGAN* to the petal lobe stands in contrast to its more restricted role as a nectar guide anthocyanin activator in other *Mimulus* species. Transcriptome sequencing of a *MYB5a* RNAi line reveals the degree to which other regulators of the anthocyanin pathway - including R3 MYB repressors and bHLH and WD40 co-activators - are responsive to the level of expression of *MYB5a*. Overall, this work reveals that a genetically simple change, which we hypothesize to be a regulatory mutation in *cis* to *MYB5a*, has cascading effects on gene expression, not only on the genes downstream of *MYB5a* but also on all of its known partners in the anthocyanin regulatory network.

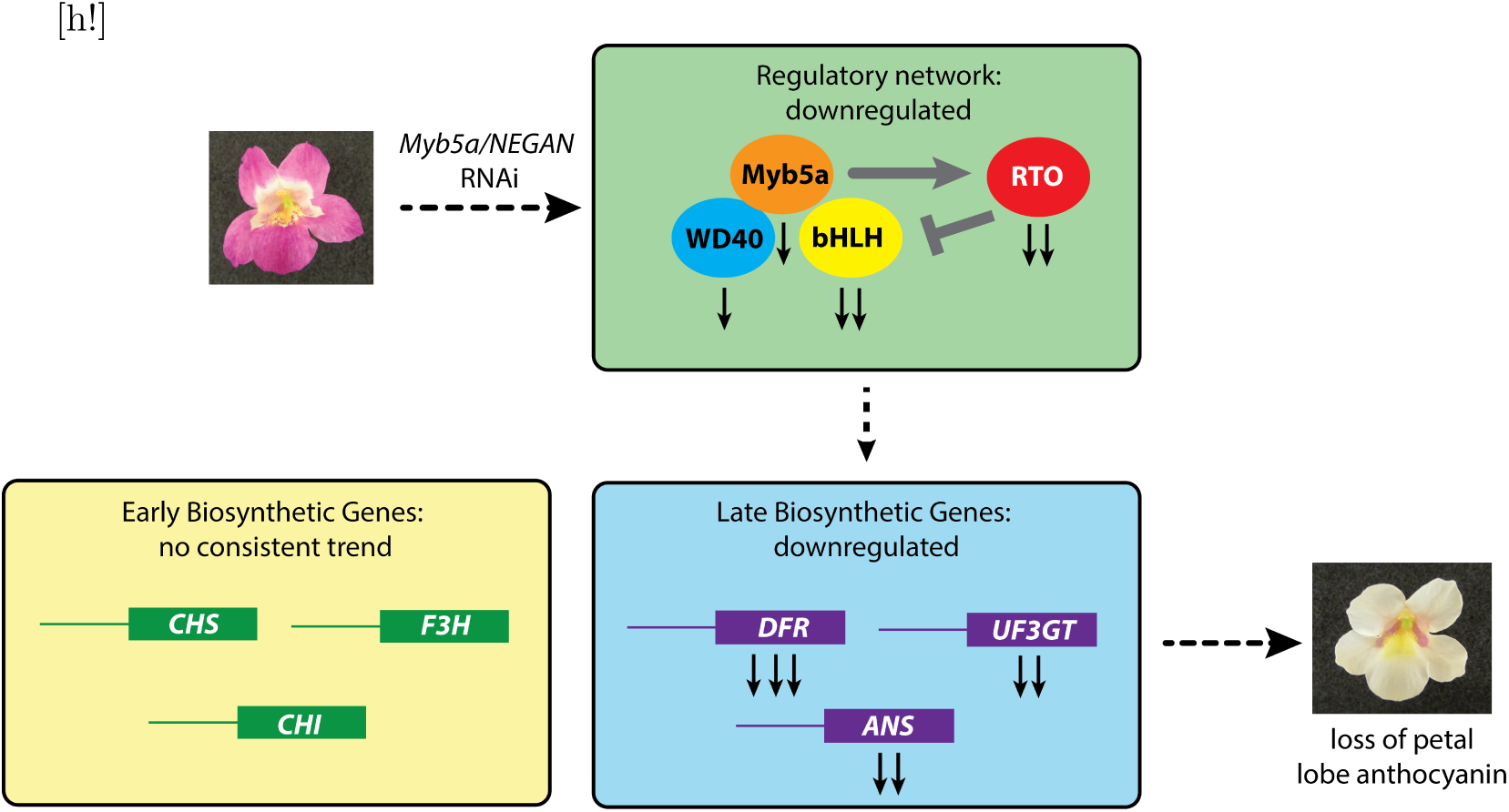
Graphical abstract. Solid black arrows indicate the direction (though not magnitude) of gene expression change, following RNAi knockdown of *MYB5a/NEGAN* in *M. l. variegatus*. The number of black arrows corresponds to the number of gene copies identified in the transcriptome. Grey symbols denote positive and negative regulatory interactions. RTO is an R3 MYB protein that inhibits anthocyanin biosynthesis by sequestering bHLH proteins away from the MBW complex.

## Introduction

Anthocyanins, the red and purple pigments that color many plant tissues, are largely responsible for the tremendous visual diversity of floral pigmentation in angiosperms. The evolutionary lability of anthocyanin pigmentation, combined with its ecological and agricultural importance, have led to the development of anthocyanin biosynthesis and regulation as a model for investigating the genetic mechanisms for trait evolution.

In dicots, the anthocyanin biosynthetic pathway is functionally divided into the Early Biosynthetic Genes (EBGs), which are activated in *Arabidopsis* by Subgroup 7 R2R3 MYB transcription factors [1], and the Late Biosynthetic Genes (LBGs), which are activated by an MBW complex composed of a Subgroup 6 R2R3 MYB (M) transcription factor and bHLH (B) and WD40 (W) co-factors [2, 3, 4, 5, 6]. Regulation of MBW is itself multifaceted. The complex is hypothesized to exhibit multiple autoregulatory feedback loops, and is counteracted by an R3 MYB which inhibits pigment production by competing with the R2R3 MYB for bHLH cofactors [7, 8, 9].

Evolutionary changes in the type of anthocyanin produced tend to occur primarily through mutations in enzyme-encoding genes [10]. In contrast, changes in pigment intensity, abundance, or location tend to evolve via the transcription factor loci - and disproportionately often in the R2R3 MYB genes compared to the bHLH or WD40 genes [10]. It is unclear whether this pattern arises because of relatively limited pleiotropy of the R2R3 MYB genes, their ability to coordinately affect expression of the late biosynthetic genes, or other properties of the network. Identifying the specific genes and molecular mechanisms responsible for pigment evolution across diverse taxa will assist in understanding these evolutionary trends.

The monkeyflower genus *Mimulus* has emerged as a useful system for investigating the evolution and regulatory mechanisms of anthocyanin regulation [11, 12, 13]. This genus was recently split into *Mimulus* and *Erythranthe* [14], but much of the monkeyflower community has opted to keep the *Mimulus* designation [15]. In species as distantly related as *M. lewisii* and *M. guttatus* - approximately 20 MY divergent [16] - the R2R3 MYB gene *MYB5/NEGAN* appears to be a key activator of anthocyanin spots within the nectar guide region of the corolla [17, 18]. A paralogous gene, *MaMYB2/PELAN*, is responsible for the solid anthocyanin pigmentation of petals in both *M. lewisii* and *M. aurantiacus* [19, 17], which diverged approximately 30 MYA ago [16]. Although petal anthocyanin has been gained and lost repeatedly across the genus [20], too few species have been studied to determine whether the spatial partitioning between *MYB5/NEGAN* and *MaMYB2/PELAN* is a consistent feature of monkeyflower evolution.

Within the *luteus* group of *Mimulus* from Chile [21, 22, 23, 24], an expansion of floral anthocyanin pigmentation into the petal lobes has evolved repeatedly, and has been genetically mapped in each case to one of two clusters of tandemly arrayed R2R3 MYB genes [25]. The *pla1* locus, responsible for the gain of petal anthocyanin in *luteus*-group species *M. cupreus* and *M. naiandinus*, includes a cluster of MYB genes that are orthologous to *MaMyb2/PELAN*. This finding is consistent with the regulatory model for petal pigmentation in *M. aurantiacus* and *M. lewisii*. In *M. luteus* var. variegatus, however, the gain of petal anthocyanin maps to the genetically unlinked locus *pla2*, containing genes *MYB4* and *MYB5* [25].

Here we use sequence analysis, quantitative and qualitative RT-PCR, transgenic manipulation of gene expression, and transcriptome analysis to investigate the gain of petal anthocyanin pigmentation in *M. l. variegatus*. We identify *MYB5a*, homologous to the nectar-guide pigmentation gene *NEGAN* from *M. lewisii* [17], as the gene responsible for this evolutionarily recent trait. Transgenic experiments reveal that the spatial domain of *MYB5a* spans both the nectar guide and petal lobe regions of the corolla: downregulating the gene via RNAi eliminates both nectar guide spots and the solid pigmentation in the petal lobes, while overexpressing the gene in the yellow-flowered *M. l. luteus* results in increased nectar guide spotting as well as petal anthocyanin pigmentation. Transcriptome analysis allows us to quantify the downregulation of *MYB5a* that was achieved in our RNAi line, and identifies a corresponding downregulation of two homeologous anthocyanin-related bHLH genes, one anthocyanin-related WD40 gene, and two homeologs of the R3 MYB repressor *Red Tongue* (*RTO*), as well as the late biosynthetic genes *DRF, ANS*, and *UF3GT*.

The results highlight how a network, and its two resulting phenotypes of nectar guide and petal lobe pigmentation, can respond in a dramatic and coordinated fashion to a relatively modest change in the expression of a single component part. They also provide a counter-example to previous work in *Mimulus*, which has documented that anthocyanin activation in the nectar guide spots versus petal lobes is partitioned between two anciently diverged paralogs.

## Materials and methods

### Plant sources and growth conditions

Seeds were originally collected from natural populations in central Chile in 2004, and were inbred for multiple generations prior to the work described here (Table 1). Seeds were sown in 2-in pots onto wet Black Gold potting soil (SunGro Horticulture, Agawam, MA) or Miracle-Gro potting soil (Scotts Miracle-Gro Co., Marysville, OH), and were misted daily until germination. For transgenic experiments, seedlings were transplanted into 6” pots to promote large size. Plants were maintained in the Whitman College greenhouse under 16-hour days and were watered daily by an automatic misting system. After expansion of the first true leaves, fertilizer (Grow Big, FoxFarm Soil Fertilizer Co., Arcata, CA, N:P:K = 6:4:4; Open Sesame, FoxFarm Soil Fertilizer Co., Arcata, CA, N:P:K = 5:45:19; and/or Miracle-Gro Bloom Booster, Scotts Miracle-Gro Co., Marysville, OH, N:P:K = 1:3:2) was applied two to three times weekly to promote growth and flowering.

**Table 1.** Seed sources.

| Taxon | Line ID | Generations Inbred | Population | Location |
| --- | --- | --- | --- | --- |
| *M. luteus* var. *luteus* | MIl-EY7 | 12 | El Yeso | 33.4°S, 70.0°W (2600 m) |
| *M. luteus* var. *variegatus* | MIv-RC6 | 11 | Río Cipreses | 34.2°S, 70.3°W (1200 m) |

Locations from which seeds were collected is given as latitude, longitude (meters above sea level).

### Expression analyses of candidate anthocyanin-activating genes

Two candidate anthocyanin-activating genes, *MYB4* and *MYB5*, had been previously identified in *M. l. variegatus* by genetic mapping and sequence characterization [25]. Because members of the *luteus* complex are allotetraploids [26, 27, 28], many genes have a homeologous copy located on a different chromosome, which could inflate expression estimates for a target gene if care is not taken to distinguish between the copies. In the process of investigating *MYB5* in the sequenced *M. l. luteus* genome, we discovered its apparent homeolog, *MYB5b* (Fig. 1A). *MYB5* will therefore be referred to in this work as *MYB5a*, to distinguish it from its homeolog. The three exons that make up *MYB5b* are identical in their coding sequence to the first three exons of *MYB5a*. However, substantial divergence in the 5’UTR (as well as downstream of the third exon) enabled the development of copy-specific primers, allowing us to disentangle the expression of *MYB5a* from its homeolog. We were not able to identify any homeolog of *MYB4* in the *M. l. luteus* genome.

**Fig 1.**
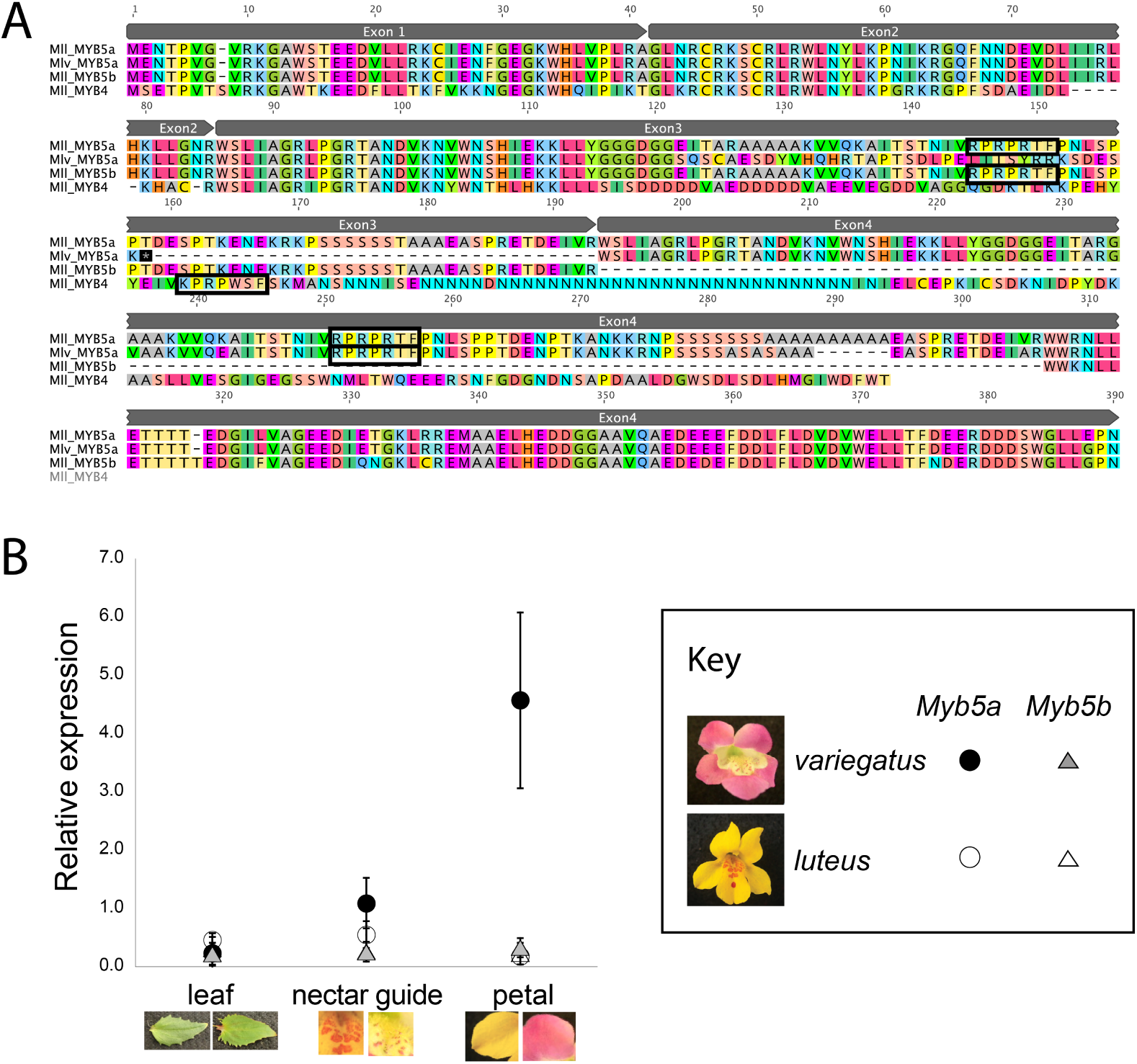
Sequence and expression of *MYB5a/NEGAN*. A. Alignment of *MYB5a, MYB5b*, and *MYB4*. Amino acid sequences were inferred by translating DNA sequences, obtained from the *M. l. luteus* genome and transcriptome [28, 31] and from PCR and Sanger sequencing of *M. l. variegatus*. Black rectangles identify putative Subgroup 6 motifs. Note that this motif, which strongly predicts ability to activate anthocyanin production, is missing from *M. l. variegatus* exon 3. Grey boxes denote exon boundaries for MYB5a and MYB5b. Created in Geneious 9.1 by Biomatters (www.geneious.com). B. qRT-PCR estimates of *MYB5a/NEGAN* and *MYB5b* expression in *M. l. variegatus* and *M. l. luteus*. Each point indicates the mean of 3 technical replicates, +/-95%CI, relative to the reference gene *Actin* and a reference pool of cDNA.

The gene responsible for activating anthocyanin biosynthesis in *M. l. variegatus* petal lobes is expected to be expressed in developing floral buds, during or just prior to the accumulation of visible anthocyanin pigment. Developmentally, this occurs approximately 72 hours before anthesis [25], before the bud has emerged from the calyx. Endpoint and quantitative RT-PCR were used to investigate patterns of expression for each candidate gene, as well as for *MYB5b*, in developing floral buds and also young leaves of the purple-lobed *M. l. variegatus* compared to the yellow-lobed *M. l. luteus*. Primers were designed based on the sequenced genome of *M. l. luteus* (Table S1), and their functionality in *M. l. variegatus* was confirmed by PCR prior to their use in expression analyses.

Although *M. l. variegatus* and *M. l. luteus* have striking differences in petal lobe pigmentation, with abundant anthocyanin production in the former and none in the latter, they share the trait of anthocyanin spotting in the nectar guide region of the corolla. Petals of developing flower buds were therefore dissected into lobe versus nectar guide components and the two tissue types were analyzed separately.

RNA was extracted using the Agilent Plant RNA Isolation Kit (Santa Clara, CA, USA). cDNA was synthesized using the ProtoScript First Strand cDNA Synthesis Kit from New England BioLabs, Inc. (Ipswich, MA, USA). Endpoint (qualitative) RT-PCR was performed on cDNA, with gDNA template as a control for primer efficacy. In addition to the *MYB* primers, either *Actin* or *GAPDH* primers were included as a control for successful cDNA synthesis (Table S1). PCRs were performed with 0.2 mM dNTPs, 0.2 uM of each primer, 0.2 uL of G-Biosciences Taq DNA Polymerase per 25-uL reaction, and 1x G-Biosciences buffer. Amplification was performed with temperature settings of 95C for 3 min, followed by thirty cycles of (95C for 30s, variable temperature for 30s, 72C for 1 minute per kb of product), and then 72C for 10 min. The annealing temperature was set to 2C lower than the lowest melting temperature of the set of primers used in the PCR, and the minimum extension time used was 30s.

Quantitative RT-PCR was performed on *MYB5a* and *MYB5b* using SYBR-Green Brilliant III Ultra-Fast reagents (Agilent Technologies, USA), in optical 96-well plates (Greiner Bio-One, Belgium), on an MxPro3000p analyzer (Stratagene, USA). Four biological replicates were used for each taxon and tissue type, collected from different plants of the same inbred line. Each biological replicate was run in triplicate and the average of the three technical replicates was used for statistical analyses. Samples were amplified for 40 cycles of 95°C for 10 s and 51°C for 20 s. Melt curves were obtained by by heating from 51°C to 95°C with a ramp speed of 0.01°C per second. The *Actin* gene was used as a normalizer. Raw qPCR fluorescence data were collected and analyzed by the default settings of the MxPro software v.4.10 (Agilent Technologies, USA). Amplification efficiencies for each primer pair were determined using the Ct (threshold) values obtained from a 1/4 dilution series (1:4, 1:16, 1:64, 1:256, and 1:1024).

### Cloning and sequencing *MYB5a* splice variants

Two splice variants of *M. l. variegatus MYB5a*, containing exons 1-2-3 and 1-2-4 respectively, were PCR-amplified using primers MYB5a-10F and MYB5a-53R (Table S1). PCR products were purified using a Genomic DNA Clean and Concentrate Kit (Zymo Research, Irvine, CA, USA), then cloned using the pGEM T-Easy kit and manufacturer’s protocol (Promega, Madison, WI, USA). pGEM vector containing the desired PCR product was transformed into competent JM109 *E.coli* cells (Promega, Madison, WI, USA). Colonies were evaluated using blue-white screening and PCR with M13 universal primers. Inserts of the expected length were sent to Eton Bioscience (San Diego, CA, USA) for Sanger sequencing.

### Transgene construction

To build transgenes using the Gateway method [29, 30], each fragment of interest was PCR-amplified using a forward primer with a 5’-CACC tag. The tag permits the fragment to be directionally cloned into a pENTR entry vector (Thermo Fisher Scientific, Waltham, MA, USA); attL/attR recombination is used to transfer the correctly oriented fragment into a Gateway-compatible destination vector.

Genomic DNA for PCR was extracted using the E.Z.N.A. SP Plant DNA Kit (Omega Bio-Tek, Norcross, GA, USA). PCR amplification was performed using Phusion High Fidelity DNA Polymerase (New England Biolabs).

Two RNAi constructs were built, targeting the 5’UTR and fourth exon respectively of *M. l. variegatus MYB5a*, with pB7GWIWG2(I) (VIB, Ghent, Belgium) as the destination vector. The inserts and primers used for transgene construction are summarized in Table 2. A previously-built overexpression construct [17], containing the complete *NEGAN* coding sequence from *M. lewisii*, driven by a 35S promoter, was also utilized.

**Table 2.**
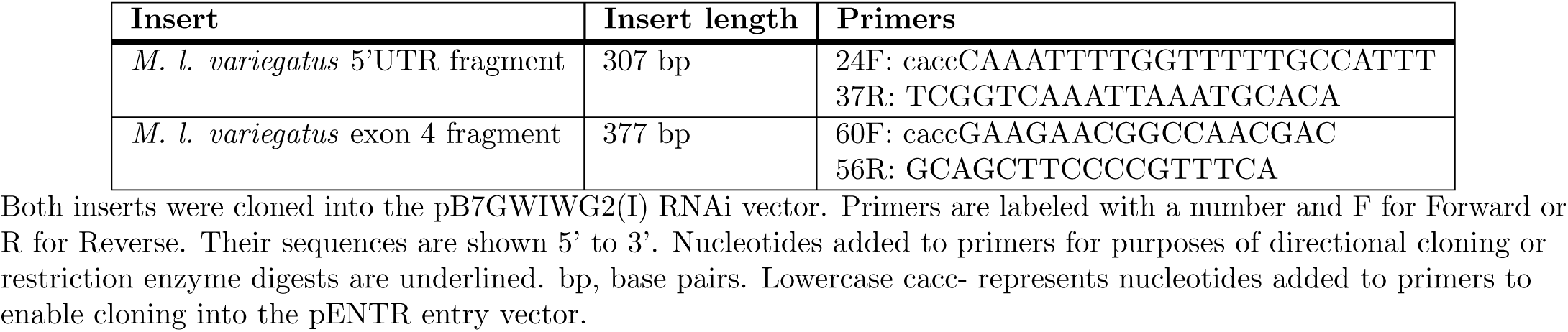
Construction of *MYB5a/NEGAN* transgenes.

### Stable transformation

Agrobacterium-mediated plant transformation was performed using floral spray and vacuum infiltration as described in [7]. Each experiment utilized approximately 12 robustly budding two- to six-month-old plants growing in 6” pots. The commercial *Agrobacterium tumefaciens* strain LBA4404 (ThermoFisher Scientific) produced only two transformed seedlings from the infiltration of 67 plants (K. Om, senior thesis 2017). Subsequent experiments were performed using strain GV3101, and this yielded higher rates of transformed seedlings.

The concentration of Silwet-L77 (Lehle Seeds, Round Rock TX, USA) used by Yuan et al. [7] was 0.1%, or 1 mL per L of *Agrobacterium* culture. This unusually high concentration of Silwet was previously found to increase transformation success in *Mimulus* (Y.-W. Yuan, unpubl. data). Acetosyringone was added to a final concentration of 0.1 M as in [7].

The predominantly-outcrossing taxa *M. l. luteus* and *M. l. variegatus* were manually self-pollinated for two weeks after the first post-infiltration flower opened. Seeds were collected and densely sown in 25 x 50 –cm flats. As soon as germination was observed, flats were sprayed daily with 1:1000 Finale (Farnam Companies, Inc., Phoenix, AZ) to eliminate non-transgenic plants. Transgenic seedlings were grown to flowering and photographed. gDNA was extracted and the presence of the transgene was verified by PCR, using a forward primer targeted to the transgene and a reverse primer targeted to the attR2 region of the transgenic plasmid (Table 2, Table S1).

RNA extractions, cDNA synthesis, and endpoint RT-PCR were performed as described in the Expression Analyses section, to evaluate the expression of the target gene relative to an untransformed control plant. Two different primer pairs targeting *MYB5a* were used, 33F-40R and 44F-45R, along with primers to reference gene *GAPDH* [31] as a positive control (Table S1).

### RNA extraction and library preparation for RNAseq

Petal tissue for transcriptome analysis was collected separately from three individuals of our highly inbred wild-type line of *M. l. variegatus*, RC6, and from three white-flowered offspring (Vrnai1.1, 1.3, and 1.5; Fig. S2) of the white-flowered RNAi transformant Vrnai1. For comparative purposes, one sample from *M. naiandinus* developing petals was also included.

Because the expression of anthocyanin-producing genes in *Mimulus* flowers is highest early in bud development, just before and after the first appearance of visible anthocyanin pigment [25, 17], we used young buds that had not yet emerged from the calyx. Anthers were removed and the remaining petal tissue (including both lobe and nectar guide regions) was snap frozen in liquid nitrogen. RNA was extracted from each of the six samples using the Agilent Plant RNA Isolation Kit (Wilmington, DE, USA). A stranded RNA-Seq plus Ribo-Zero library preparation, followed by one lane of Illumina HiSeq 4000 50-bp single-read sequencing, was performed by the Duke University sequencing core facility (Durham, NC).

### Transcriptome alignment

With a published *Mimulus luteus* var. *luteus* genome and gene feature annotation file available [28], we chose the genome splice-aware mapping approach to assemble the *M. luteus* var. *variegatus* transcriptomes to the *M. l. luteus* genome [32, 33]. Sequencing quality control was performed by plotting the sequence nucleotide distribution and sequencing quality scores for all samples. The transcriptome libraries were aligned using Bowtie2(Version 2.3.5.1) under-very-sensitive-local mode [34] for best results in distinguishing homeolog expression, given that the *M. luteus*-group species are putative tetraploids [26, 35]. Overall alignment rates were greater than 90% for all 7 samples. From the sequence alignment maps we counted reads per gene for all samples using the exon coordinates included in the published luteus GFF (general feature format) file. Read count was performed with software HT-seq (Version 0.11.2) [36] in Python. For detailed alignment documents see the supplemental data. Pipeline code, alignment documents and results and are accessible through the GitHub repository (https://github.com/cici-xingyu-zheng/Luteus-RNA-seq) and for raw read count see Table S2.

### Transcriptome analysis and functional annotation

Analyses of differential gene expression were conducted in DESeq2 (Version 1.26.0) in the R/Bioconductor environment [37](R Version 3.6.2; Bioconductor Release 3.10). After normalizing each gene by sequencing depth, we performed a principal component analysis (PCA) plot of the seven transcriptome samples (three wild-type *M. l. variegatus*, three *M. l. variegatus* from RNAi line Vrnai1, and *M. naiandinus* as an outgroup). As expected, the samples clustered by treatment with the outgroup being an outlier (Fig. S2).

After performing a shrinkage estimation for dispersion to address the inaccuracy introduced by the small sample size and reduce the false positive rate [37], the logarithmic fold change (LFC) between RNAi treatment and control samples was used to evaluate differential expression. Following a false discovery rate control using the Benjamini-Hochberg Correction method, transcripts that were log-2-fold up or down-regulated with a p-Value *<* .05 were considered to be significantly differentially expressed. Transcript expression profiles were normalized to Reads Per Kilobase Per Million (RPKM) for further analysis and for plotting (Table S3).

To annotate differentially expressed genes, all of the *M. l. luteus* gene sequences were translated to protein sequences using EMBOSS(6.5.7) “transeq” command and searched against *Arabidopsis thaliana* (TAIR) protein database (www.arabidopsis.org) using the “BLASTp” query with a e-value cut-off *<* 10^*-*6^. For each coding sequences for *M. l. luteus* were released by Edger et al. [26] along with the draft genome and can be accessed at https://datadryad.org/stash/dataset/doi:10.5061/dryad.d4vr0. For each coding sequence from the *M. l. luteus* genome, the best-hit *A. thaliana* gene was used to annotate the transcript with a gene name and Gene Ontology (GO) annotation terms.

These annotations were then applied to the *M. l. variegatus* transcriptome data. GO enrichment analysis for was conducted, for genes that were differentially expressed between wild-type *M. l. variegatus* and the *MYB5a* RNAi line Vrnai1, using topGO (Version 2.36.0) with the “org.At.tair.db” database in R. Pathway enrichment tests were done using the KEGG (the Kyoto Encyclopedia of Genes and Genomes, https://www.genome.jp/kegg/;Release 93.0) pathway assignments for Arabadopsis with the KEGGREST Bioconductor package [38].

### Regulatory and pathway analysis

In order to test for an effect of *MYB5a* downregulation on the anthocyanin biosynthetic pathway, we examined the expression of genes corresponding to six core pathway enzymes that produce the cyanidin pigment found in *M. l. variegatus* [21]. These are: chalcone synthase (CHS), chalcone isomerase (CHI), flavonoid-3-hydroxylase (F3H), dihydroflavonol-4-reductase (DFR), anthocyanidin synthase (ANS), and UDP-flavonoid-3-glucosyl-transferase (UF3GT) [39, 40]. A list of *M. l. variegatus* transcripts with *RPKM >* 1 and annotated with descriptions matching these enzymatic activities was obtained. Genes were considered truly orthogonal if they were the best hit in a reciprocal BLAST or had the same high BLAST score as a putative homeolog.

In addition to the enzyme-encoding genes, we were also interested in what other regulatory factors might be affected when we knock down the *MYB5a* transcriptional activator. In all species that have been examined, the regulatory complexes of the anthocyanin pathway include members of the MYB, basic helix–loop–helix (bHLH), and WD40 repeat families [41, 42]. Therefore, we subset from the differentially expressed genes a list of genes annotated as members of these three transcriptional regulator families. We searched within this subset for homologs to genes in *M. guttatus* and *M. lewisii* that have been reported to be involved in anthocyanin regulation.

### Data availability

Seeds, bacterial strains, and plasmids are available upon request. Supplemental files are stored in bioRxiv, DOI 10.1101/2020.04.09.030536. Individual *MYB4*, *MYB5a* and *MYB5b* coding sequences are located in GenBank. Whole-genome sequence for *M. l. luteus* was released by Edger et al. [28] along with the draft genome and can be accessed at https://datadryad.org/stash/dataset/doi:10.5061/dryad.d4vr0. Raw transcriptome data for *M. l. luteus* are published in [31] and can be accessed at http://dx.doi.org/10.5061/dryad.84655. The *M. l. variegatus* transcriptome raw reads will be made available on the Sequence Read Archive (https://www.ncbi.nlm.nih.gov/sra; link to data will be provided once upload is complete). Transcriptome analysis code and alignment documents are accessible through the GitHub repository (https://github.com/cici-xingyu-zheng/Luteus-RNA-seq).

## Results and Discussion

### Expression of an alternately spliced R2R3 MYB, *MYB5a/NEGAN*, covaries with petal anthocyanin

In *M. l. variegatus*, only two anthocyanin-related genes were found within the genetically mapped petal anthocyanin locus [25]. The two genes, *MYB4* and *MYB5a*, both grouped phylogenetically with the anthocyanin-activating Subgroup 6 of the R2R3 MYB gene family [43, 25]. Reciprocal BLAST searches of the *M. lewisii* genome indicate that *MYB5a* is the homolog of *NEGAN*, a gene shown to be responsible for nectar guide anthocyanin spots in *Mimulus lewisii* [17].

The Subgroup 6 R2R3 MYBs generally have three exons. The diagnostic Subgroup 6 amino acid motif is KPRPR[S/T]F in *Arabidopsis*, and it is located in the third exon [43]. In both *M. l. luteus* and *M. l. variegatus*, however, MYB5a genomic sequence data revealed the presence of a fourth exon that is extremely similar in sequence to the third exon. In both taxa, this fourth exon appears to be intact and potentially functional, including the Subgroup 6 motif. The third exon, although apparently intact in *M. l. luteus*, has a 26-bp frameshift-inducing deletion in *M. l. variegatus* that eliminates the critical Subgroup 6 domain and creates a premature stop codon (Fig. 1A).

*MYB4*, while somewhat similar in sequence to *MYB5a/NEGAN*, has a nonconservative amino acid substitution in the functionally critical DNA-binding motif that is a hallmark of Subgroup 6 transcription factors (Fig. 1A) and has a top BLAST hit on NCBI to *M. guttatus* LOC105953416, which encodes a GL1-like trichome differentiation protein. Additionally, unsuccessful efforts to amplify *MYB4* out of *M. l. variegatus* floral bud cDNA, using primers that successfully amplify the gene from gDNA, indicate that this gene is not detectably expressed in developing *M. l. variegatus* buds (Fig. S1). Consistent with its similarity to a trichome differentiation gene, we did detect expression of *MYB4* in leaf tissue (Fig. S1).

In contrast, quantitative RT-PCR revealed that *MYB5a/NEGAN* is strongly and specifically expressed in the anthocyanin-pigmented petal lobes of *M. l. variegatus*, while showing little to no expression in the non-anthocyanin-pigmented petal lobes of *M. l. luteus* (Fig. 1B). Expression of *MYB5a/NEGAN* was modest in the anthocyanin-pigmented nectar guide tissue of both taxa, and undetectable in their leaf tissue (Fig. 1B). The homeologous gene copy, *MYB5b*, showed consistently low expression across both taxa and all tissue types, at levels less than 10% of that observed in *M. l. variegatus* petal lobes (Fig. 1B).

Two alternate splice variants of *MYB5a* were discovered, via RT-PCR followed by cloning and Sanger sequencing. An “exon 1-2-3” transcript was occasionally isolated from developing *M. l. variegatus* petals, but was never found in *M. l. luteus*, even in the anthocyanin-pigmented nectar guides. An “exon 1-2-4” transcript was reliably isolated from *M. l. variegatus* petal lobes and from the spotted nectar guide tissue of the corolla throat. In *M. l. luteus*, the exon 1-2-4 transcript was recovered from nectar guides but not from the (anthocyanin-free) petal lobe tissue. Thus, the exon 1-2-4 transcript of *MYB5a* appears to be consistently expressed in all anthocyanin-pigmented floral tissue, and absent from tissue that lacks anthocyanin.

### *MYB5a/NEGAN* is sufficient and necessary for petal anthocyanin in *M. l. variegatus*

A pilot experiment showed that *NEGAN* from *M. lewisii*, when overexpressed with the 35S promoter, was sufficient to activate anthocyanin production in both leaf and petal tissue of the normally yellow-flowered *M. l. luteus* (Fig. 2A). Anthocyanin pigmentation in the nectar guide region was also dramatically increased relative to wild-type (Fig. 2A). Encouraged by this result, we used a 35S-driven RNAi transgene to test the hypothesis that the exon 1-2-4 transcript of *MYB5a* is necessary for petal lobe anthocyanin pigmentation in *M. l. variegatus*. In the strongest line, “Vrnai1,” anthocyanin pigmentation was completely abolished in both the petal lobe and the nectar guide (Fig. 2B). Anthocyanin pigmentation in the stems, the leaves, and the adaxial side of the corolla appeared to be unaffected, indicating that the effect of *MYB5a* knockdown is spatially specific. Successful knockdown of *MYB5a* in line Vrnai1 was demonstrated via qualitative RT-PCR (Fig. 2C), and confirmed through transcriptome analysis (next section).

**Fig 2.**
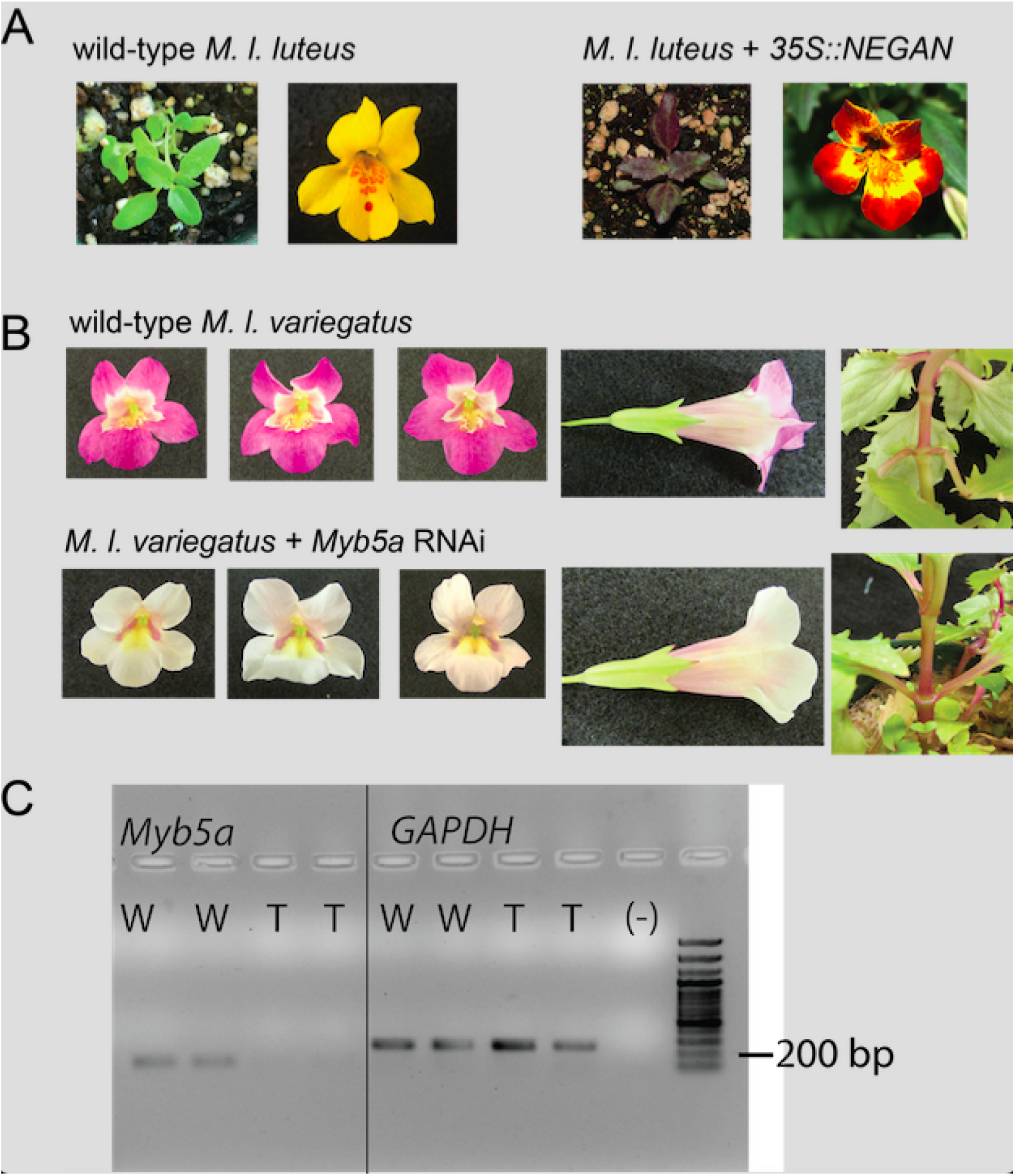
*MYB5a/NEGAN* is sufficient and necessary for activation of petal lobe anthocyanins. A. Overexpression of the coding sequence of *MYB5a/NEGAN* from *M. lewisii* in the normally yellow-petaled *M. l. luteus* activates anthocyanin biosynthesis in both leaf and petal tissue. B. RNAi targeting *MYB5a* exon 4 in the normally purple-petaled *M. l. variegatus* eliminates anthocyanin biosynthesis in the petal lobes and nectar guides, but not elsewhere in the plant. The strongest of 8 RNAi lines, Vrnai1, is shown here. See Supplemental Data for images of the other RNAi lines. C. Qualitative (end-point) RT-PCR on cDNA from developing petal lobes of wild-type (W) and transgenic (T) *M. l. variegatus* reveals a reduction in *MYB5a* expression in the Vrnai1 transgenic line. Reference gene *GAPDH* was used as a positive control.

Considerable variation in the severity of the RNAi phenotype was observed across different transgenic lines, as is typical for this technique (Fig. 3). From 12 wild-type *M. l. variegatus* transformed with RNAi targeting exon 4 of *MYB5a*, six putatively transgenic seedlings were obtained. Four of these (Vrnai1, Vrnai3, Vrnai5, Vrnai7) had a visible reduction in petal lobe anthocyanin pigmentation. From an additional 12 wild-type *M. l. variegatus* transformed with RNAi targeting the 5’UTR of *MYB5a*, two putatively transgenic seedlings were obtained. One of these (Vrnai6) had a visible reduction in petal lobe anthocyanin pigmentation. Genotyping confirmed the presence of the RNAi transgene in all eight lines (Supp Fig.s: Genotyping gels). In line Vrnai6, the RNAi phenotype declined over the lifetime of the plant (Fig. 3). This phenomenon has been shown to be caused by methylation and epigenetic silencing, particularly of the 35S promoter, in both *Arabidopsis thaliana* and the tobacco species *Nicotiana attenuata* [44, 45, 46].

**Fig 3.**
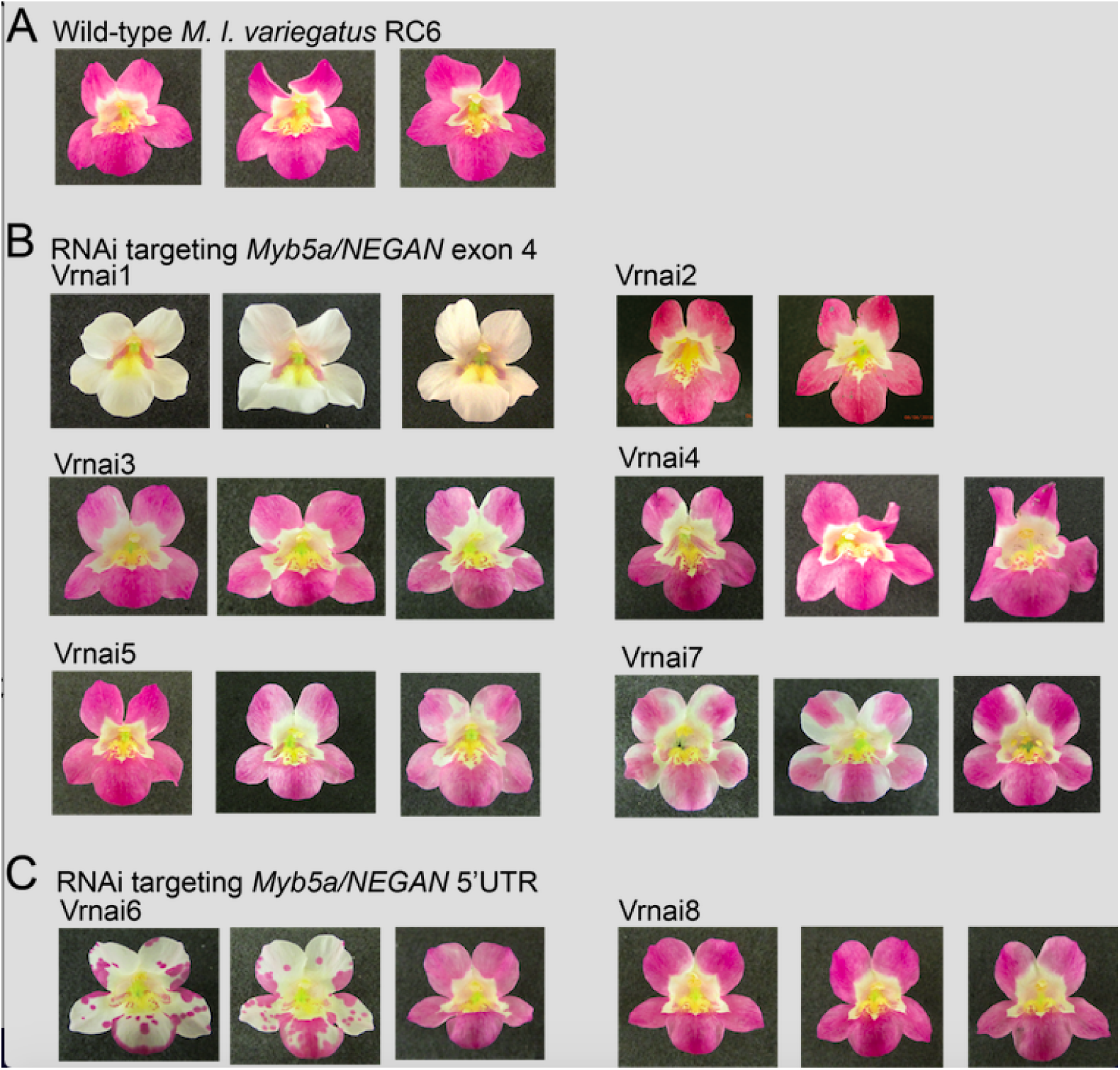
Transgene effects vary between and within insertion lines. A. Wild-type *M. l. variegatus*. B. Stable transgenic lines, for which the presence of an RNAi transgene targeting *MYB5a/NEGAN* exon 4 was confirmed by genotyping. C. Stable transgenic lines, for which the presence of an RNAi transgene targeting *MYB5a/NEGAN* 5’UTR was confirmed by genotyping.

The Vrnai1 plant was self-fertilized, and eight of the resulting seeds were planted and grown to flowering. As expected for a single-copy (heterozygous) insertion of a dominantly-acting transgene in the Vrnai1 plant, we observed a 3:1 ratio of six white-flowered and two wild-type plants among the offspring (Fig. S2). The six white-flowered offspring exhibited slight anthocyanin mottling on some flowers (Fig. S2).

### Transcriptomic differences between wild-type and RNAi lines of *M. l. variegatus*

We found a total of 632 genes that were significantly differentially expressed between wild-type and Vrnai1 lines of *M. l. variegatus*, with 346 genes down-regulated and 290 up-regulated (Table S4). The differentially expressed genes are enriched in a variety of functions including response to UV-B, anthocyanin-containing compound biosynthesis, pollen exine formation and phenylpropanoid biosynthetic process (p-value *<* .001) (Table S5). A pathway enrichment analysis identified two significantly enriched pathways (Table S6): (1) cutin, suberine and wax biosynthesis, and (2) flavonoid biosynthesis, which includes the anthocyanin biosynthetic pathway [39].

The *M. l. variegatus MYB5a* DNA sequence, which had previously been determined by PCR and Sanger sequencing, had best hits to three coding sequences in *M. l. luteus*: Mlu 12200, Mlu 12207, and Mlu 42095. In the *M. l. variegatus* transcriptomes, the latter two transcripts were not expressed at all in either wild-type or RNAi lines of *M. l. variegatus*. In contrast, Mlu 12200 was robustly expressed in all six libraries, with three-fold higher expression in the wild-type compared to the *MYB5a* RNAi line (Fig. 4A). The Mlu 12200 transcript from *M. l. variegatus* also had a best match to the gene that is annotated as *MYB5a* in the *M. l. luteus* genome, [25, 28], further confirming its identity as the target of our RNAi experiment.

**Fig 4.**
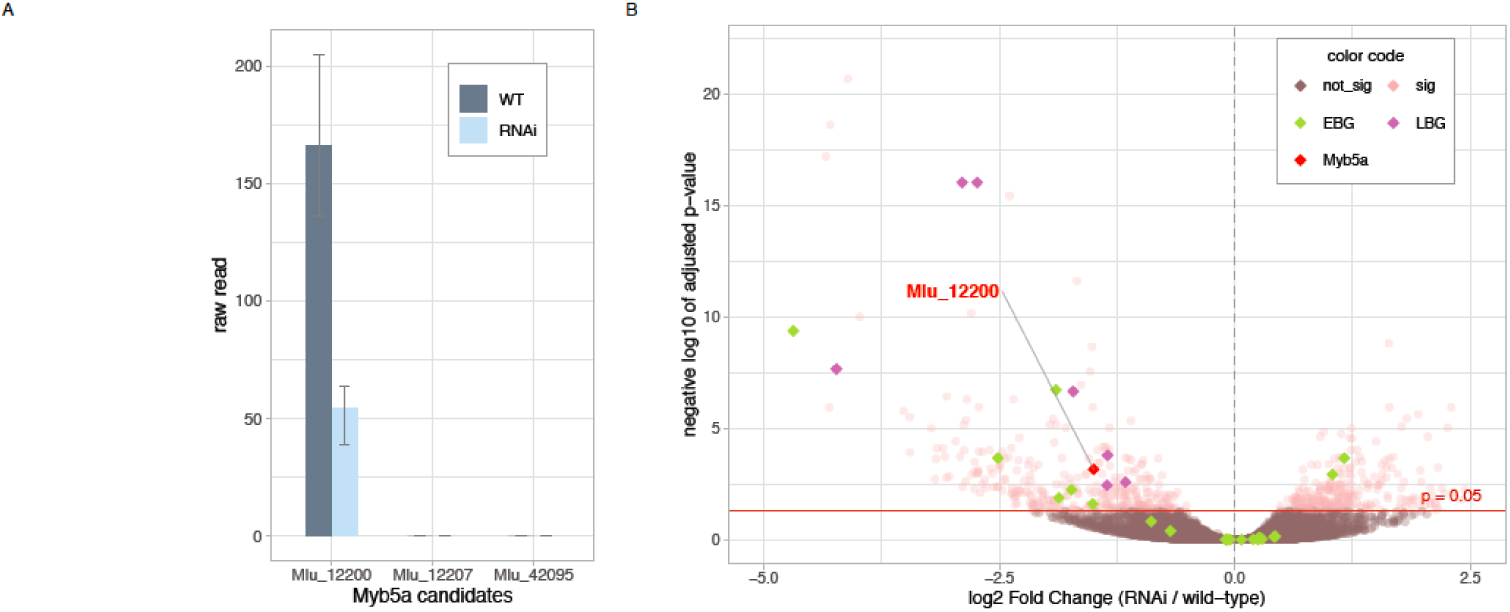
RNAi knockdown of *MYB5a* (transcript Mlu 12200) results in coordinated downregulation of the Late Biosynthetic Genes in the anthocyanin biosynthesis pathway. A. Experimental confirmation of the correspondence of transcript Mlu 12200 to *MYB5a*. Bars show the average read-count (n=3) of wild-type and RNAi lines respectively, for candidate transcripts Mlu 12200, Mlu 12207, and Mlu 42095 from the RNA-seq Transcriptome Libraries. Upper and lower error bars mark the highest and lowest read counts; only Mlu 12200 had any mapped reads. B. Differentially expressed anthocyanin biosynthesis genes. Log-2 fold change in expression (Vrnai1 / wild-type) is shown on the x-axis, and negative log 10 of adjusted p-value in the y-axis. 30 structural genes were identified, with Early Biosynthetic Genes (EBGs) shown in green and Late Biosynthetic Genes (LBGs) in purple. Points in the upper left quadrant correspond to genes that are significantly down-regulated in the RNAi line relative to wild-type *M. l. variegatus*. sig, significant expression difference between Vrnai1 and wild-type; not sig, no significant difference.

To check for off-target effects of the RNAi transgene, we asked whether any other Subgroup 6 R2R3 MYB genes were significantly down-regulated in the RNAi line. Five MYBs were identified among the down-regulated genes (Mlu 24690, Mlu 05348, Mlu 27563, Mlu 17841, and Mlu 00921), but none of them contained a Subgroup 6 motif. We conclude that the loss-of-pigment phenotype observed in line Vrnai1 is due to the reduction in *MYB5a* expression alone, and was not caused by incidental down-regulation of another anthocyanin-activating MYB gene.

### RNAi knockdown of *MYB5a* reduces transcription of the late anthocyanin biosynthesis genes

We identified a total of 30 genes that are annotated to have enzymatic functions corresponding to the six core enzymes in the cyanidin pathway [39] (Table S7). The Early Biosynthetic Genes (*CHS, CHI* and *F3H*) are particularly enriched in copy number, each having more than the two homeologous copies expected in a tetraploid.

In eudicots, the anthocyanin biosynthetic genes are usually divided into two groups, the early (EBGs) and late (LBGs) biosynthetic genes, with the latter being tightly regulated by an MBW complex [1, 2, 3, 4, 5, 6]. In Arabidopsis, *CHS, CHI*, and *F3H* belong to the EBGs, producing the precursors of not only flavonoid pigments but other flavonol compounds. *DFR, ANS*, and *UF3GT* comprise the LBGs. Conserved as the enzymatic pathway is, however, the break point of regulation between early and late genes can vary across species [47]. We utilized our transcriptomic data to determine where the breakpoint occurs in *M. l. variegatus*.

Consistent with expectations from the literature, the anthocyanin biosynthetic genes in *M. l. variegatus* show a dichotomous pattern of response to *MYB5a* down-regulation. The genes earlier in the pathway - *CHS, CHI*, and *F3H* - had no consistent pattern of expression change (Fig. 4B, S4), while the genes later in the pathway - *DFR, ANS*, and *UF3GT* - were consistently down-regulated (Fig. 4B, S5). All copies of the LBGs had statistically significant change in the same direction (lower expression in the *MYB5a* RNAi line relative to wild-type), while the multiple copies of the EBGs showed a mix of expression increase, decrease, and no change in the RNAi line. Our data suggest that *MYB5a* controls anthocyanin production by regulating primarily the late biosynthetic genes, which include *DFR, ANS*, and *UF3GT*. This conclusion is consistent with the proposed mechanism of anthocyanin pathway regulation in the congeneric *M. aurantiacus* [19].

### RNAi knockdown of *MYB5a* reduces transcription of other anthocyanin regulators

Within the list of differentially expressed genes, we identified 20 genes that are annotated to be transcription factors with MYB or helix–loop–helix (HLH) domains, or that encode a WD40 protein that could potentially belong to a MYB–bHLH–WD40 regulatory complex (Table S8). Since different MBW complexes regulate a variety of traits besides anthocyanin synthesis, we further examined these candidates to identify the most likely homologs of anthocyanin-specific MBW components. BLAST results indicate that one WD40 gene (labeled *MlutWD40a*) and two bHLH genes (*MlutbHLH1* and *MlutbHLH2*) have high sequence similarity to anthocyanin *TTG1* and *TT8* genes, which are the main WD40 and bHLH regulators respectively of proanthocyanidin biosynthesis in Arabidopsis seed [6]. *MlutWD40a* appears to be an ortholog of *MlWD40a* from *M. lewisii*, which has been shown to co-activate anthocyanin expression in the corolla [17]. Interestingly, the two bHLH co-factors have the highest similarity to *M. lewisii ANbHLH3*. In *M. lewisii*, this gene is not detectably expressed in the petal lobes [17], suggesting a functional diversification in the bHLH co-factors as well as in *MYB5a/NEGAN*.

Recent studies show that single-repeat R3 MYBs are activated by the R2R3 MYB member of the MBW complex, and inhibit anthocyanin biosynthesis by directly interacting with the bHLH component of the same complex [48, 7]. We found that two R3 MYB genes, Mlu 13044 (*MlutR3MYB1*) and Mlu 33990 (*MlutR3MYB2*), have high sequence similarity to *RTO* from *M. lewisii* [49] and are strongly and significantly down-regulated in the *MYB5a* RNAi line. Their downregulation in Vrnai1 is consistent with the model that *MYB5a* is an activator of its own inhibitor, although the impact of *MYB5a* on two apparently homeologous gene copies is unique.

We were particularly interested in the expression change of these five anthocyanin related regulators when knocking down *MYB5a*. In previous studies, the R2R3 MYB protein has been shown to activate the complex as well as the R3 inhibitor, and we hypothesized that in *M. l. variegatus* RNAi lines, the other candidate anthocyanin regulators we identified would show the same directional expression change as *MYB5a* [50]. *Indeed, our expression results showed that down regulating MYB5a* in the RNAi line of *M. l. variegatus* led to a down regulation in all the putative regulatory genes, highlighting the critical role that *MYB5a* plays in the MBW regulatory complex (Fig. 5). The activators of anthocyanin biosynthesis - including R2R3 MYB, bHLH, and WD40 transcription factors - were all 2- to 3-fold down-regulated in the RNAi line (Fig. 5). The two R3 MYB inhibitors showed much more dramatic effects: they were 19- and 29-fold down-regulated (Fig. 5). High sensitivity of the R3 MYB inhibitors to *MYB5a* expression may be a key feature of the anthocyanin regulatory network.

**Fig 5.**
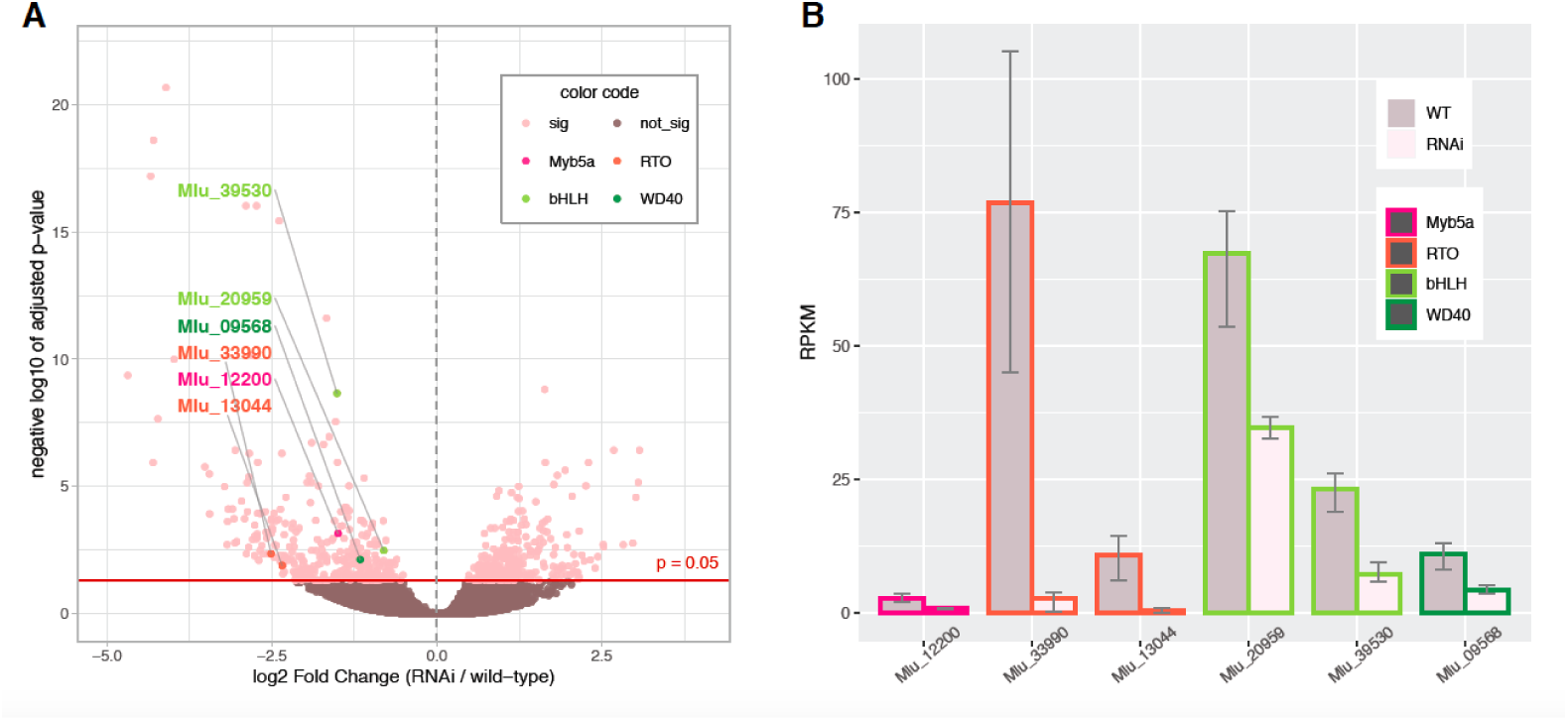
RNAi knockdown of *MYB5a* results in downregulation of multiple other anthocyanin regulatory genes. A. Expression of the regulatory complex genes relative to the transcriptome as a whole. Log-2 fold change in expression (Vrnai1 / wild-type) is shown on the x-axis and negative log 10 of adjusted p-value as the y-axis. Points in the upper left quadrant correspond to genes that are significantly down-regulated in the RNAi line relative to wild-type *M. l. variegatus*. Genes with homology to known anthocyanin-regulating genes are color-coded as *MYB5a* (pink); the R3 MYB repressor *RTO* (orange); bHLH (light green); or WD40 (dark green). sig, significant expression difference between Vrnai1 and wild-type; not sig, no significant difference. B. Transcript level fold change of the anthocyanin-regulating transcription factors. Normalized expression fold change of the regulatory genes is shown in RPKM (per million mapped reads). Upper and lower error bars represent maximum and minimum expression level among the samples (n=3).

## Conclusions

Here we demonstrate that *MYB5a* is both sufficient and necessary for the evolutionarily recent gain of petal lobe anthocyanin in the purple-flowered *Mimulus luteus* var. *variegatus*. We identify an unusual four-exon structure to this Subgroup 6 R2R3 MYB, along with evidence for alternative splicing. While the role of coding versus *cis*-regulatory evolution was not explicitly examined in this paper, we found that patterns of expression of *MYB5a* corresponded to pigmentation: the gene was strongly expressed in the heavily anthocyanin-pigmented petal lobes of *M. l. variegatus*; not detectably expressed in the petal lobes of *M. l. luteus*, a conspecific which lacks petal lobe anthocyanin; and modestly expressed in the partly-pigmented nectar guides of both taxa. Further supporting the importance of expression change rather than coding change is that the homologous coding sequence from the evolutionarily distant *M. lewisii*, which is approximately 20 MY divergent from *M. l. variegatus* [16], was highly effective at activating anthocyanin in the normally yellow-flowered *M. l. luteus.* Additional transgenic experiments, such as promoter swaps, will be helpful in testing the impact of coding versus *cis*-regulatory evolution on the functional divergence of *MYB5a* in *M. l. variegatus*.

Our transgenic experiments indicate that *MYB5a* is responsible for anthocyanin pigmentation in two distinct regions of the *M. l. variegatus* corolla: the petal lobe and the nectar guide. The role of *MYB5a/NEGAN* in nectar guide spotting appears to be conserved across much of the genus. In contrast, the deployment of *MYB5a/NEGAN* to the petal lobes has not, to our knowledge, been previously reported. Rather, ancient paralogs of *MYB5a/NEGAN* appear to be responsible for petal lobe anthocyanin in diverse species including *M. lewisii, M. aurantiacus, M. cupreus*, and *M. naiandinus* [25, 17, 19, 51].

Transcriptome analyses of an RNAi line of *M. l. variegatus* confirmed the targetted knockdown of *MYB5a*, and highlighted the inter-relatedness of expression patterns across the network of anthocyanin regulatory and biosynthetic genes. These results illustrate how the network can respond dynamically to expression changes of a single network component, creating an avenue for a relatively modest expression change – such as that caused by a single *cis*-regulatory mutation - to have a major effect on the transcriptome as well as on the ultimate phenotype.

## Supporting information

**S1 Fig.**
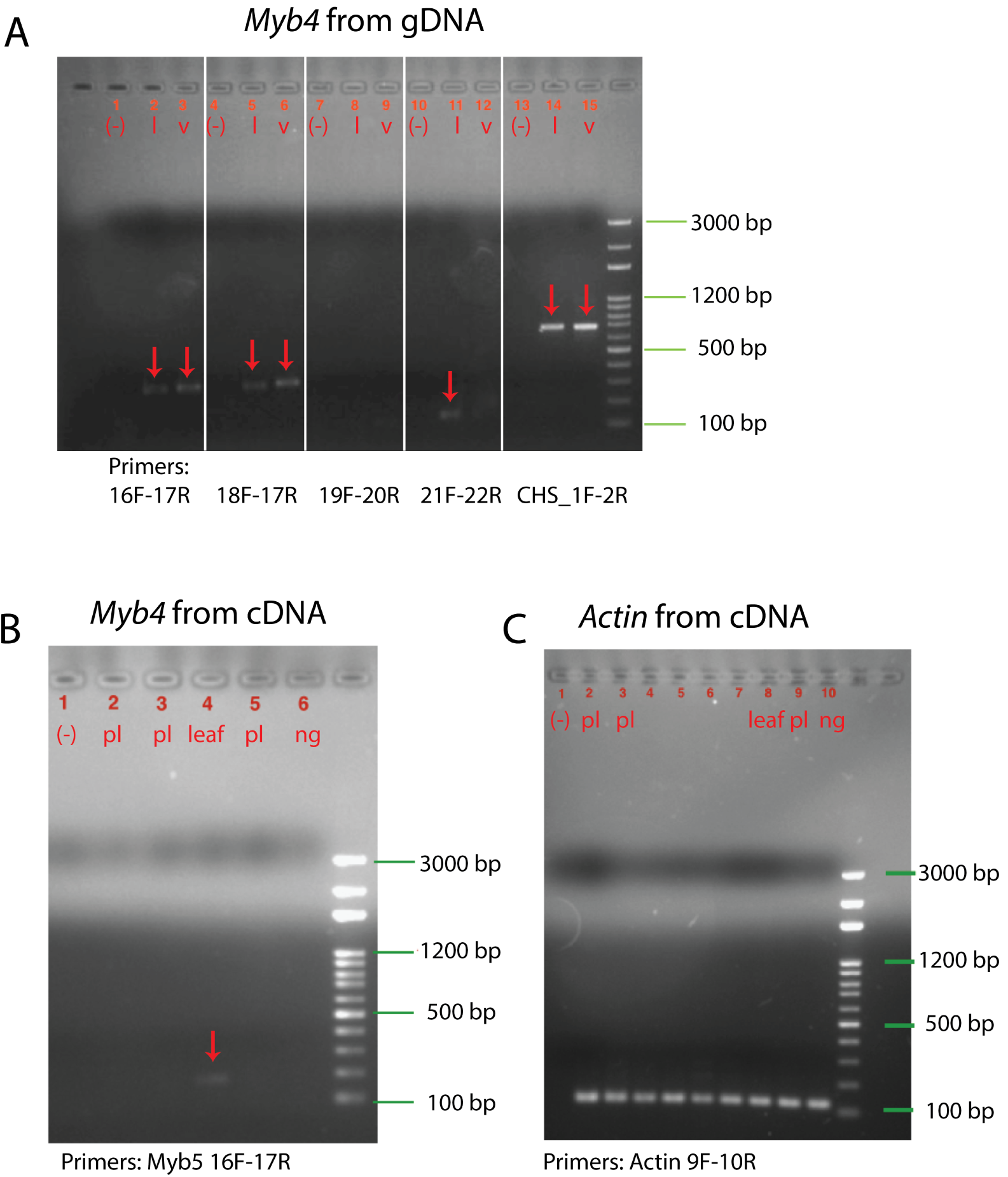
RT-PCR indicates a lack of *MYB4* expression in developing *M. l. variegatus* petal lobes. A. *MYB4* primers 16F-17R and 18F-17R successfully amplify a *MYB4* fragment from *M. l. luteus* and *M. l. variegatus* genomic DNA. Expected lengths are 310 base pairs and 320 base pairs, respectively. (-), no-template control; l, *M. l. luteus*; v, *M. l. variegatus*. The *CHS* gene is included as a positive control; expected length is 650 base pairs. B. The *MYB4* 16F-17R primers amplify a *MYB4* fragment out of *M. l. variegatus* leaf cDNA, but not out of developing *M. l. variegatus* floral bud cDNA. Expected length from cDNA, for this intron-spanning primer pair, is 190 base pairs. (-), no-template control; pl, petal lobe; ng, nectar guide. C. The *Actin* reference gene was used as a positive control to verify cDNA quality. (-), no-template control. The cDNA samples used in panel C are labeled as in panel B: pl (petal), leaf, or ng (nectar guide).

**S2 Fig.**
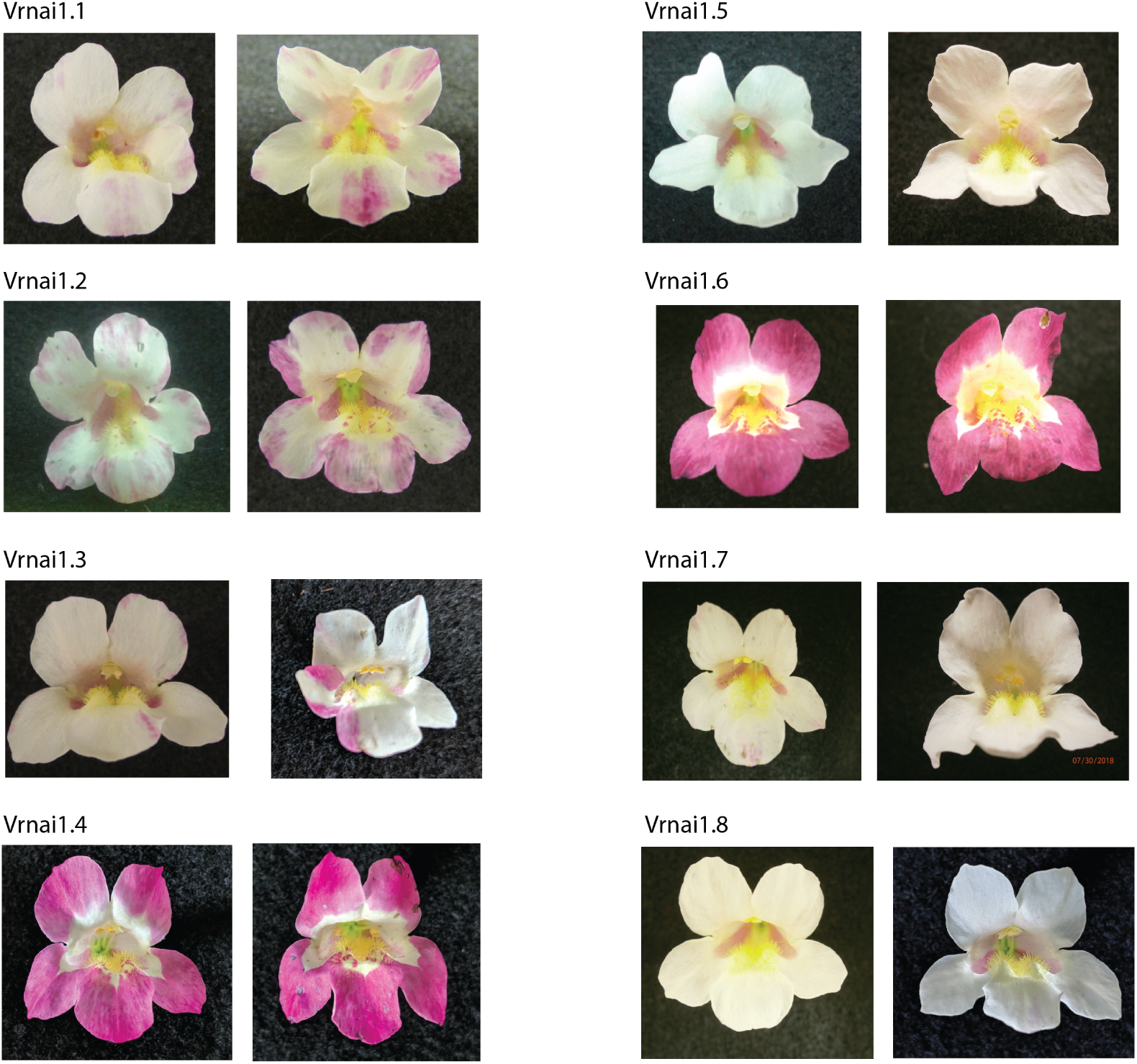
Offspring of line Vrnai1 show a 3:1 ratio of RNAi phenotype : wild-type, consistent with a single transgene insertion in Vrnai1. Eight seedlings were grown to flowering. Two flowers per plant are shown. Plants 1.1, 1.3, and 1.5 were used for RNA extraction and transcriptome sequencing.

**S3 Fig.**
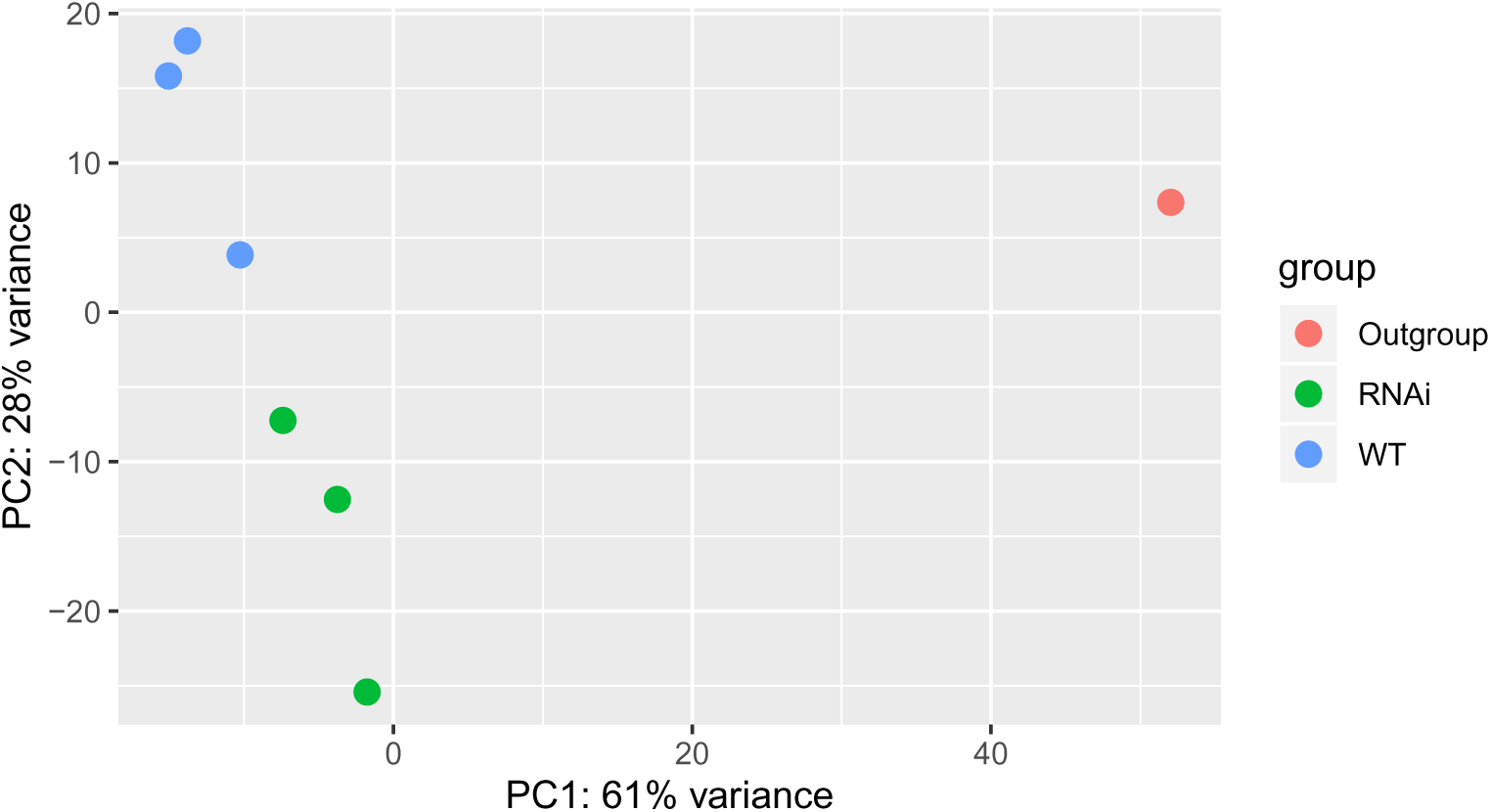
Principle components analysis illustrating transcriptome differences between *MYB5a* RNAi and wild-type *M. l. variegatus*. Unsupervised clustering of the 7 transcriptome libraries was performed using principle component analysis based on the differences between samples in normalized read count of all genes. The x and y axes represent the 1st and 2nd principal components.

**S4 Fig.**
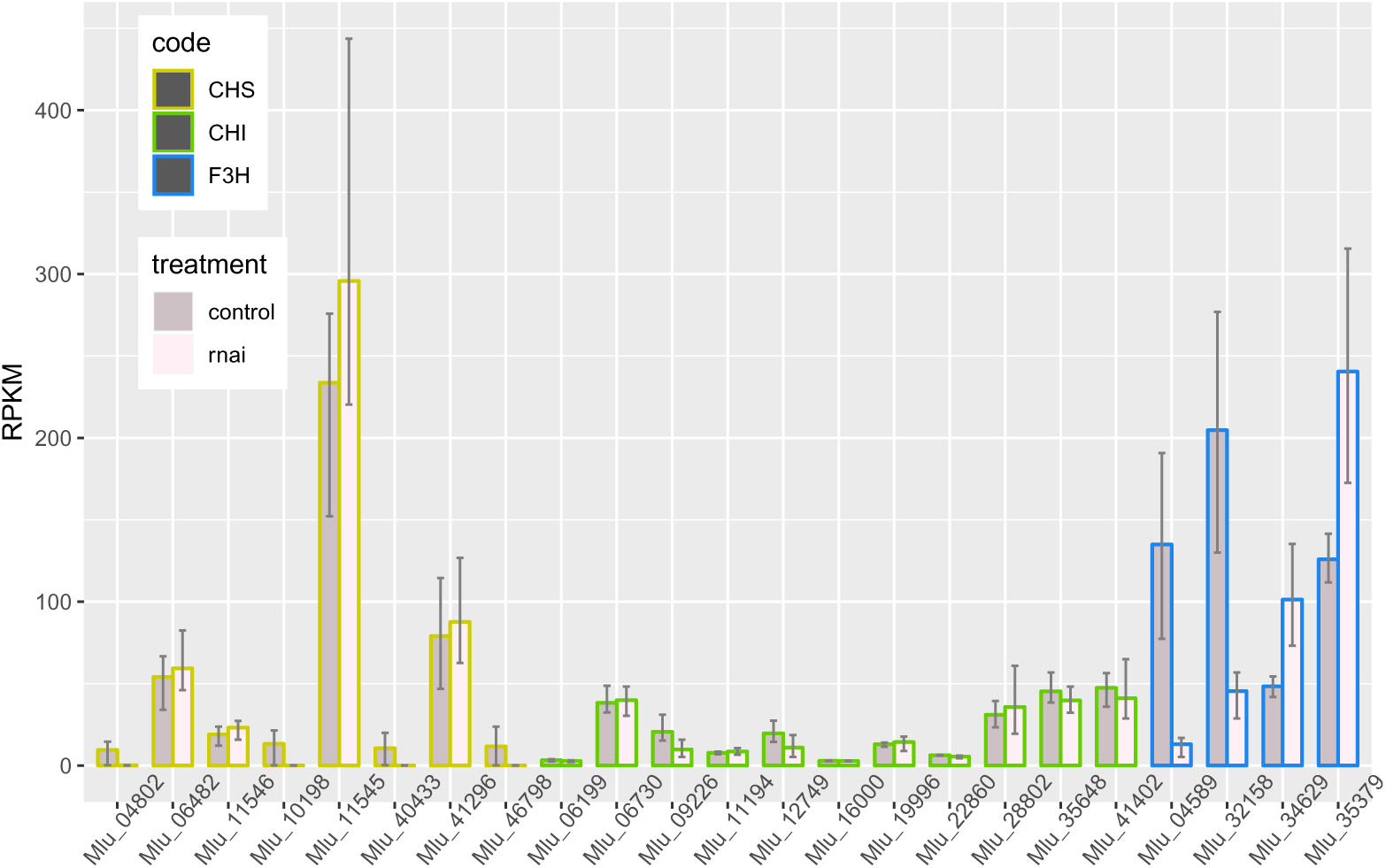
Transcript level of Early Biosynthetic Genes (EBGs) of the anthocyanin pathway in *MYB5a* RNAi line Vrnai1 compared to wild-type *M. l. variegatus*. No consistent pattern in expression level is observed comparing the RNAi lines to the wild-type. The bars represent the average expression level, and upper and lower error bars represent maximum and minimum expression level among the samples (n=3). RPKM (per million mapped reads) is used as the normalized unit of transcript expression.

**S5 Fig.**
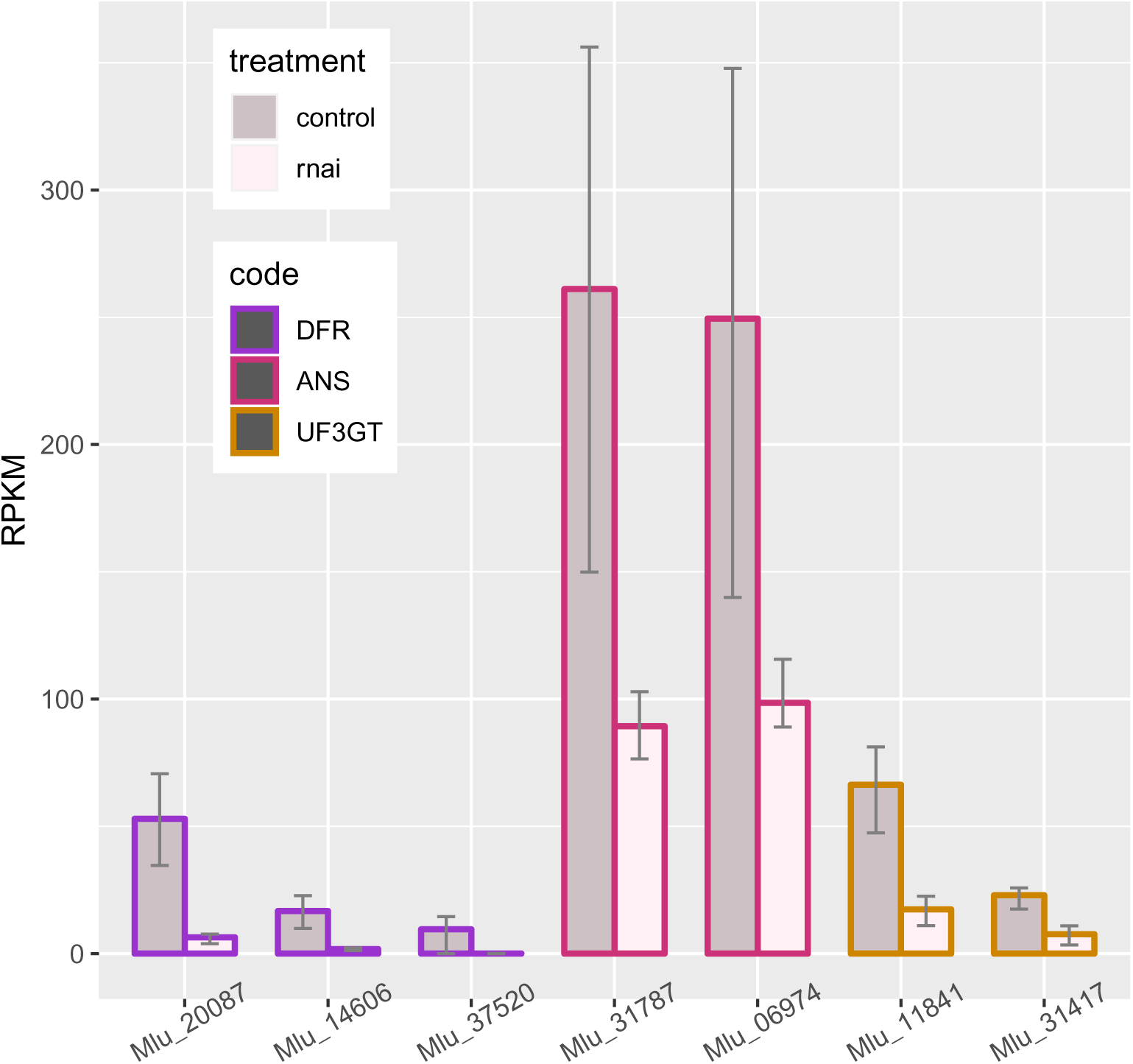
Transcript level of Late Biosynthetic Genes (LBGs) of the anthocyanin pathway in *MYB5a* RNAi line Vrnai1 compared to wild-type *M. l. variegatus*. All genes identified as LBGs show a decreased expression in the RNAi line compared to the wild type. The bars represent the average expression level, and upper and lower error bars represent maximum and minimum expression level among the samples (n=3). RPKM (per million mapped reads) is used as the normalized unit of transcript expression.

**S1 Table. Primers used for qualitative and quantitative RT-PCR and transgene construction.** Primers are named by their target gene(s), an identification number, and “F” for forward primers or “R” for reverse primers.

**S2 Table. Raw read count of all transcriptome libraries from HTSeq.** S1 indicates the outgroup, *M. naiandinus*. S2-S4 are from the RNAi line Vrnai1, and S5-S7 are from wild-type *M. l. variegatus*.

**S3 Table. Normalized transcript expression level of all libraries in RPKM.** Mlv-wt, wild-type [untransformed] *M. l. variegatus* line RC6. Vrnai1, *MYB5a* RNAi line 1. Mna, *M. naiandinus* line 105.

**S4 Table. Significantly differentially expressed genes from DESeq2 results.** Result of all 632 differentially expressed genes from DESeq2 analysis, giving base means across samples, log2 fold changes, standard errors, test statistics, p-values and adjusted p-values.

**S5 Table. Significance of GO terms in the differentially expressed genes.** First we did a classical enrichment analysis by testing the over-representation of GO terms within the group of differentially expressed genes (Fisher’s Test) and then performed Kolmogorov-Smirnov test using the both the “classic” and the “elim” method. S5 gives a data frame containing the top GO terms identified by the elim algorithm with p-value cut of at .01.

**S6 Table. KEGG Pathway Enrichment Testing of the differentially expressed genes.** We accessed KEGG pathway assignments for Arabidopsis through the KEGGREST Bioconductor package, and then applied Wilcoxon rank-sum test to each pathway for enrichment testing. Columns features pathway code, pathway name, the p value of being enriched and the number of annotated genes in the pathway.

**S7 Table. Forward and reciprocal BLAST results for anthocyanin pathway genes.** The column “code” contains the short-hand annotation for the structural genes of the anthocyanin pathway; the column “blastp-besthit” shows the best hit in Arabidopsis; the column “rcp-blastp-besthit” shows the reciprocal results for each best-hit Arabidopsis genes. Genes are highlighted if the reciprocal BLAST identifies the same *M. l. luteus* gene.

**S8 Table. Forward and reciprocal BLAST results for regulatory genes.** The column “code” contains the short-hand annotation for the potential regulatory genes of the anthocyanin pathway; the column “blastp-besthit” shows the best hit in Arabidopsis; the column “rcp-blastp-besthit” shows the reciprocal results for each best-hit Arabidopsis genes. Genes are highlighted if the reciprocal blast identifies the same *M. l. luteus* gene.

## Acknowledgments

The authors gratefully acknowledge assistance with molecular work from Baoqing Ding, Lauren Stanley, Rachel Eguía, Johanna Au, Jonah Rodewald, and Jeanette Schwensen. We thank Larry North for construction of a vacuum infiltration apparatus, Carter Cooke and Bella Rivera for greenhouse assistance, and Amy LaFountain and Qiaoshan Lin for assistance in obtaining transgenic seeds. This material is based upon work supported by the Murdock College Research Program for Life Sciences Award 2015270, NSF microMORPH training grant, and NSF-DEB Award 1655311, to AMC.

